# NcPath: A novel tool for visualization and enrichment analysis of human non-coding RNA and KEGG signaling pathways

**DOI:** 10.1101/2022.06.03.494777

**Authors:** Zutan Li, Yuanyuan Chen, Yuan Zhang, Jingya Fang, Zhihui Xu, Hao Zhang, Minfang Mao, Liangyun Zhang, Cong Pian

**Affiliations:** College of Science, Nanjing Agricultural University, Nanjing, 210095, China; College of Agriculture, Nanjing Agricultural University, Nanjing, 210095, China; Simcere Diagnostics Co., Ltd., Nanjing, 210023, China

**Author notes:** To whom correspondence should be addressed., Correspondence may also be addressed to Lianyun Zhang. The authors wish it to be known that, in their opinion, the first three authors should be regarded as Joint First Authors.

## Abstract

Noncoding RNAs play important roles in transcriptional processes and participate in the regulation of various biological functions, in particular miRNAs and lncRNAs. Despite their importance for several biological functions, the existing signaling pathway databases do not include information on miRNA and lncRNA. Here, we redesigned a novel pathway database named NcPath by integrating and visualizing a total of 178,308 human experimentally-validated miRNA-target interactions (MTIs), 36,537 experimentally-verified lncRNA target interactions (LTIs), and 4,879 experimentally-validated human ceRNA networks across 222 KEGG pathways (including 27 sub-categories). To expand the application potential of the redesigned NcPath database, we identified 553,523 reliable lncRNA-PCG interaction pairs by integrating co-expression relations, ceRNA relations, co-TF-binding interactions, co-Histone-modification interactions, *cis*-regulation relations and lncPro Tool predictions between lncRNAs and protein-coding genes. In addition, to determine the pathways in which miRNA/lncRNA targets are involved, we performed a KEGG enrichment analysis using an hypergeometric test. The NcPath database also provides information on MTIs/LTIs/ceRNA networks, PubMed IDs, gene annotations and the experimental verification method used. In summary, the NcPath database will serve as an important and continually updated platform that provides annotation and visualization of the pathways on which noncoding RNAs (miRNA and lncRNA) are involved, and provide support to multimodal noncoding RNAs enrichment analysis. The NcPath database is freely accessible at http://ncpath.pianlab.cn/.

## Introduction

Understanding the mechanisms of gene regulation is a major challenge in molecular biology and bioinformatics. The rapid development of high-throughput sequencing technologies [1] and the emergence of new *multiomics* technologies [2, 3] significantly expanded research on gene regulation at the transcriptional, post-transcriptional, translational and post-translational levels. Previous studies showed that RNA-protein interactions regulate gene expression by controlling various post-transcriptional processes, which in turn directly or indirectly affect disease development [4]. The dysregulation of noncoding RNAs, in particular microRNAs (miRNAs) and long noncoding RNAs (lncRNAs), is closely associated with a variety of biological processes and disease development [5], whereby it is essential to obtain additional references and evidence to elucidate the molecular mechanisms involved [6].

MicroRNAs (miRNAs) are small non-coding RNAs with around 18-26 nucleotides that are transcribed from DNA sequences into primary miRNAs, and subsequently processed into precursor and mature miRNAs in animal and plant species. Since the first miRNA gene Lin-4 was discovered in 1993[7], more than 35,000 miRNA sequences have been identified across more than 270 organisms [8]. MicroRNA can induce mRNA decapping and alkenylation via base binding with the complementary sequence of the 3′ untranslated region (3′UTR), which negatively regulates gene expression by accelerating mRNA degradation or suppressing mRNA translation, affecting major pathways in a post-transcriptional fashion [9-12]. A high number of studies reported that miRNAs play a role in crucial various cell activities, such as cell cycle [13], cell proliferation [14], differentiation [15], apoptosis [14], metabolism [16], cellular signaling [17], embryonic development [18], virus defense [19], and hematopoietic processes [20]. In addition, miRNAs have been associated with several diseases, especially in different types of cancer [21-23], whereby these non-coding RNA molecules represent good candidate biomarkers for potential diagnosis and prognosis of cancer and other diseases [24].

The other important class of pervasively noncoding RNAs, lncRNAs, constitute an heterogeneous group of RNA molecules >200 nucleotides long [25-28], which play critical roles in a wide range of biological processes and might thus be used as novel biomarkers [29-36]. Accumulating evidence suggests that lncRNAs, acting as oncogenes or tumor suppressors, play complex and precise regulatory roles in cancer initiation and progression [37, 38]. Importantly, it has also been reported that lncRNAs regulate the proliferation, differentiation, invasion, metastasis and metabolic reprogramming of cancer cells [39-41]. Furthermore, lncRNAs play important functional roles in regulating the transcription and translation of metabolism-related genes, acting as decoys, scaffolds, and competing with endogenous RNAs (ceRNAs) [42-44].

Analysis of target genes and target pathways in the context of systems biology is a crucial step for noncoding RNA research, in particular studies on disease that include comparisons between experimental groups. To this end, several analytical tools and bioinformatic databases have been developed. In the case of miRNAs, several research tools with varying scope and functionality were developed, including: MIENTURNET [45], miRNet [46] and miRViz [47], which generate miRNA-target interaction networks; miRTarVis [48] that can be used to visualize co-expression networks of paired miRNA and mRNA data; miRUPnet [49], miEAA [50], BUFET [51] and miSEA [52], which are databases providing enrichment analysis for miRNAs; the online database miRNApath [53] and the R package CORNA [54], which incorporate GO and KEGG enrichments obtained from predicted and validated miRNA-target interactions; miRTar [55], a tool that links individual miRNAs to metabolic pathways; miTALOS v2 [56], a program that has been developed to analyze miRNA functions and tissue-specific regulation in signaling pathways; DIANA-mirPath v3.0 [57], a web server used for miRNA pathway analysis that can be used to predict miRNA targets through the DIANA-microT-CDS algorithm; miRPathDB 2.0 [58], which indexes enriched pathways for known miRNAs and miRNA candidate genes using validated and predicted target genes from the literature; miRTargetLink 2.0 [59], an interactive tool for miRNA research that dynamically presents miRNA target genes and pathway networks; miRPathDB 2.0 [58] and miRTargetLink 2.0 [59], two recently published databases for which the pathway information supports interpretability and which focus on miRNA pathways, somehow limiting their scope. As for tools available for lncRNA analysis, those with an application scope closest to the redesigned NcPath proposed here include: NONCODE [60], LNCipedia [61] and RNAdb [62], which are comprehensive databases that provide basic annotation information for lncRNAs; Co-LncRNA [63], Lnc-GFP [64], LncTarD [65], LncR2metasta [66] and FARNA [67], which were developed to infer lncRNA biological functions; NPInter [68], lnCeDB [69], starBase v2.0 [70], DIANA-LncBase [71], miRSponge [72] and PceRBase [73], which provide information on lncRNA-target relationships. Of these, DIANA-LncBase [71] collects experimentally verified and predicted miRNA-lncRNA pairs; LncRNA2Target v2.0 [74], which provides predicted lncRNA-target relationships; LncReg [75], which contains experimentally-validated records of lncRNA-associated regulatory entries; and LncACTdb 3.0 [76], which supports comprehensive information on ceRNAs across different species and their involvement in different diseases.

Combining the above database of miRNAs and lncRNAs can greatly promote our understanding and propel research efforts on non-coding RNAs. However, not only the data and methods presented in most databases need to be updated, but also most of these tools fail to simultaneously integrate mRNA, miRNA and lncRNA for comprehensive analysis. In addition, only few studies explored the integration of both miRNA and lncRNA with pathway databases. The development of a more comprehensive relationship of noncoding RNAs and gene regulation in human biological pathways thus represents an urgent research goal at present. Importantly, enrichment analyses based on reference genes and noncoding RNA (miRNA and lncRNA) sets will be useful for analyzing noncoding RNA lists of interest submitted by different users.

In order to facilitate the study of pathway-related miRNA, we reported the first version of the NcPath database (miR+Pathway), which allows users to search relationship networks of 115 pathway-related miRNAs and provides an miRNA-based gene enrichment analysis tool. Since it was first released in 2019, several more miRNA-target interactions (MTIs) and lncRNA target interactions (LTIs) were identified, in particular through experimentally validated approaches. Hence, this work stems from the need to update the old version with more resources and functions to improve the tool. Accordingly, we developed a novel human pathway visualization database named NcPath (Figure 1 and Table 1), which focuses on accommodating human pathway information with various available resources for noncoding RNA-target interactions, and performs visualization and enrichment analysis of non-coding RNA lists submitted by users. NcPath supports 178,308 human experimentally-validated miRNA-target interactions (MTIs) and 36,537 experimentally-verified lncRNA target interactions (LTIs) across 222 KEGG pathways (including 5 main categories and 27 sub-categories). We also recorded 4,879 experimentally-validated human ceRNAs relationships between lncRNAs and miRNAs. Furthermore, by integrating co-TF-binding interactions, co-Histone-modification interactions, ceRNA relations, *cis*-regulation relations and lncPro Tool predictions between LncRNAs and protein-coding genes in dozens of databases, we were able to identify 553,523 reliable lncRNA-PCG interaction pairs involved in different gene pathways. Moreover, gene enrichment analysis associated with gene regulation, miRNAs and lncRNAs can be performed simultaneously.

**Table 1.**
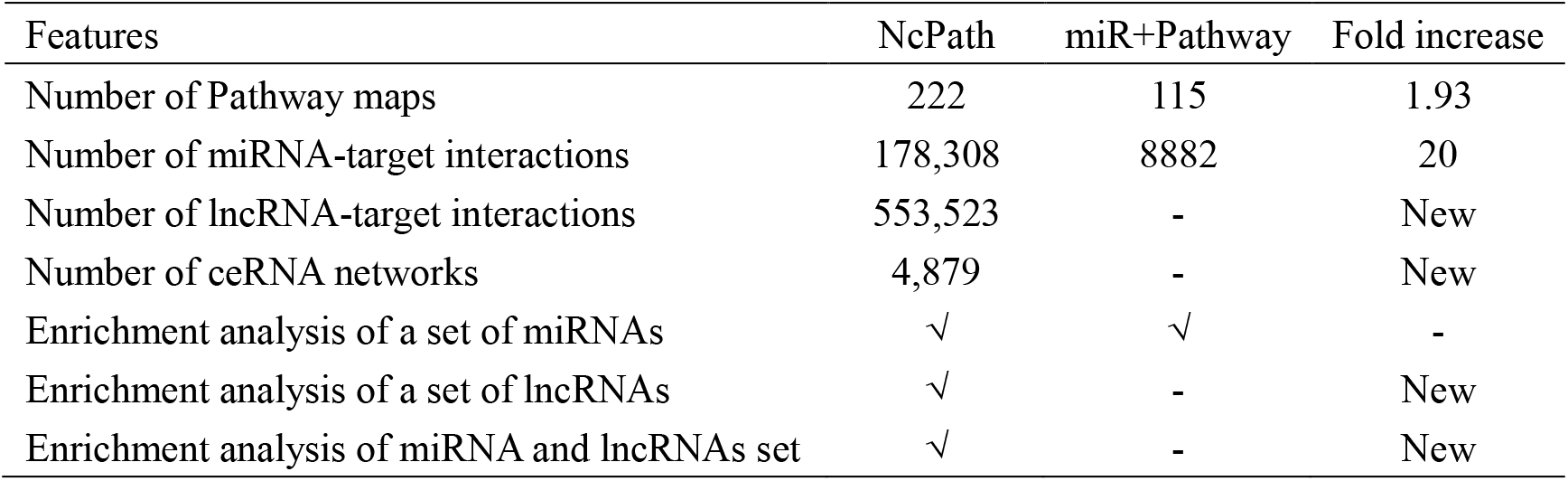
Data content and function of NcPath and comparison of the current and previous versions

**Figure 1.**
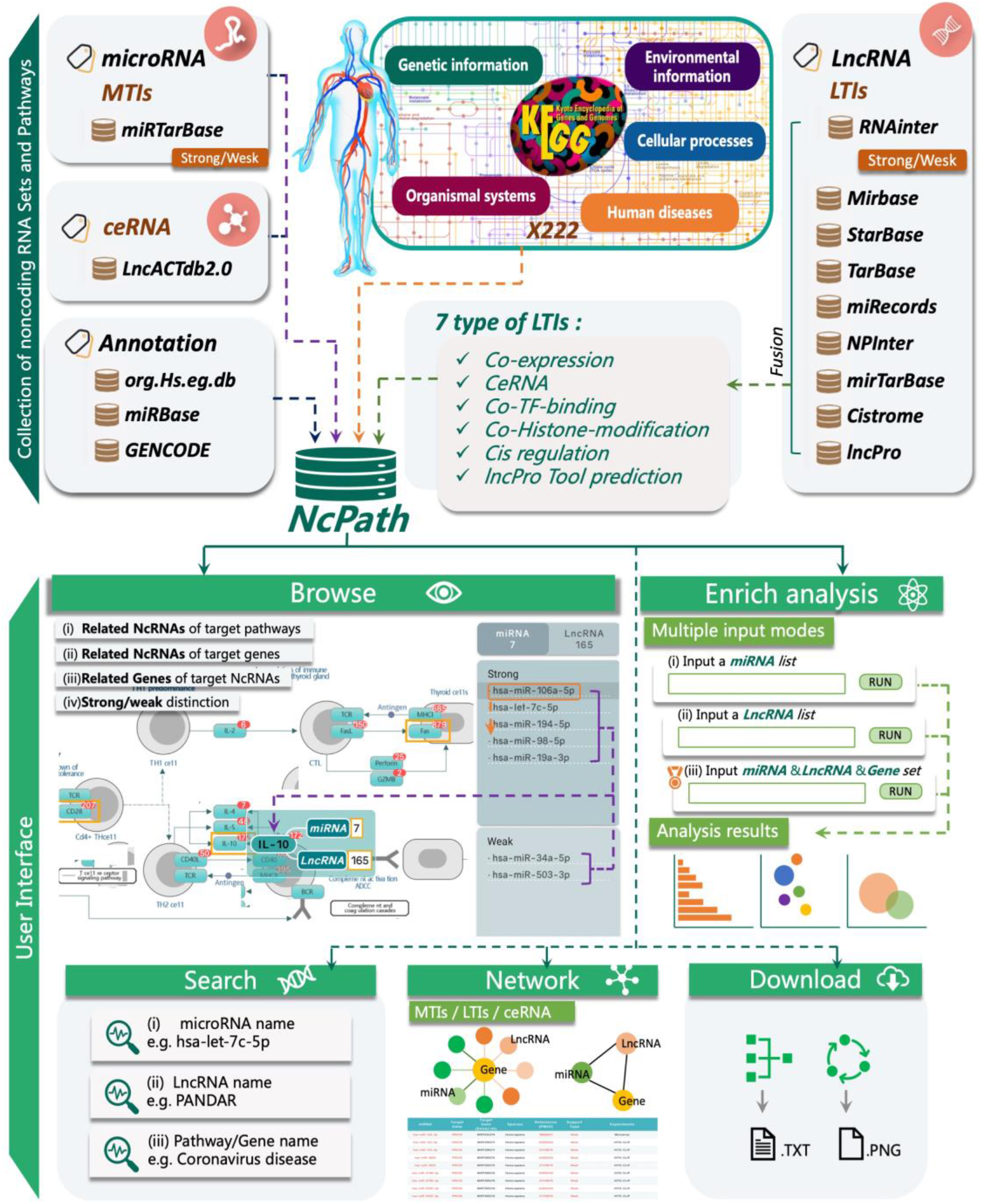
Collection of datasets and user interface of NcPath. The upper panel represents the database content that includes experimentally-validated noncoding RNAs and predicted noncoding RNAs. The lower panel refers to the user interface of NcPath, which supports multiple functions, including browsing, enrichment analysis, search, interaction networks and download. The pathway interaction visualization with noncoding RNAs is also provided in NcPath.

NcPath bridges the gap between biological pathways and human lncRNAs/miRNAs to enhance our understanding of lncRNAs/miRNAs function, particularly the investigation of the roles played by these types of RNAs in physiological and pathological processes. In summary, NcPath constitutes a more complete platform that provides two types of noncoding RNA sets in pathways for users, and performs visualization and enrichment analysis of noncoding RNA sets submitted by users.

## MATERIALS AND METHODS

### Data selection and processing

The new version of NcPath supports non-coding RNAs annotation records and targets for *Homo sapiens*, which were obtained from the latest release of org.Hs.eg.db (Release 3.11) [77], miRBase (v.22.1) [78], and GENCODE [79]. In the case of miRNAs, the validated MTIs were acquired from miRTarBase v.8 [80] which has accumulated >2 200 449 verified MTIs. The ceRNA relationships between lncRNA, miRNA and mRNA were obtained from LncACTdb3.0 [76]. For lncRNAs, the experimentally-verified lncRNA target interactions (LTIs) were collected from RNAinter [81]. We obtained seven types of reliable lncRNA-mRNA interactions, for which the specific details are as follows:

i. **Co-expression relation between LncRNAs and PCGs**. We used the annotation file of the whole transcriptome downloaded from GENCODE V33 [79] (including 22,742 protein coding genes and 17,907 lncRNAs) and expression data downloaded from GTEx (we selected 27 normal tissues with each sample size more than 30, including 7,806 samples) to compute the Pearson correlation coefficient (PCC) and adjusted P-value (fdr) between lncRNA and PCG, respectively. We then used the mean PCC of 27 tissues as the final co-expression score of lncRNA and PCG.
ii. **ceRNA relation between lncRNAs and mRNAs**. The interaction relationships of miRNAs (2,656 miRNAs in the miRBase database [78]) and their target genes were downloaded from 8 distinct databases, specifically: StarBase [70], TarBase [82], miRecords [83], NPInter [84], mirTarBase [80]; miRNA-lncRNA: StarBase [70], NPInter [84]; DIANA-LncBase [71]). We collected 99,010 miRNA-lncRNA and 1,869,961 miRNA-mRNA interaction pairs in total. An hypergeometric test was performed to identify the ceRNA relation of lncRNAs and mRNAs. The corrected BH P-value and the shared miRNAs of each lncRNA-mRNA pair were then calculated.
iii. **Co-TF-binding interaction of LncRNA-PCG pairs**. A total of 11,348 human transcription factor (TF) ChIP-seq and 11,079 human histone modification (HM) ChIP-seq datasets involving 1,117 TFs and 77 histone modification patterns, respectively, were collected from the Cistrome database [85]. The datasets with less than 30 targets were removed, with 9,489 TF ChIP-seq and 9,384 histone modification datasets left for analysis. Considering that a higher number of common TFs (HMs) combined with lncRNA and mRNA led to a stronger co-existence, we measured the possibility of co-existence using the Jaccord coefficient:

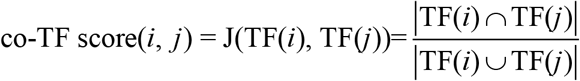

TF (*i*) and TF (*j*) represent the TF set interacting with the mRNA *i* and the lncRNA *j*, respectively.
iv. **Co-Histone-modification interaction of LncRNA-PCG pairs**. Similarly, the co-HM score was defined to measure the possibility of co-existence in terms of histone modification. This definition assumes that the pairs of identified genes were subjected to co-selection pressures (more fortuitous than expected) during evolution and are therefore considered to be functionally related.
v. **lncRNA cis regulation relation**. Genes located 20KB upstream or downstream of a lncRNA were considered *cis*-regulated by the lncRNA. LncRNA and mRNA annotation files were downloaded from GENCODE [79], with a total of 21,128 *cis*-regulated lncRNA-mRNA relationship pairs being identified.
vi. **lncRNA-protein interactions identified by Clip-seq**. We collected the human lncRNA-PCG interactions from the NPInter v4.0 database. We only retained lncRNA-PCG interactions identified by Clip-seq and that were present in the GENCODE annotation file [79] of V33. Finally, 36,363 lncRNA-PCG relationship pairs were identified by Clip-seq.
vii. **lncRNA-protein interaction pairs identified by the prediction tool lncPro**. Firstly, the lncRNA and protein sequences were downloaded from GENCODE V33 [79]. In order to improve efficiency, we modified the source code of lncPro [86] to calculate the properties of all proteins in batches, and then calculated the interaction scores between all transcripts and protein sequences. The final lncRNA-mRNA interaction score was defined as the mean of all possible interaction scores between multiple transcripts and multiple corresponding protein sequences. After this, we used lncPro to generate the scores (0-100) measuring the binding possibility between protein and lncRNA using sequence information.

For the specific process and results of screening the relationship between mRNA and lncRNA, please refer to the section ‘Screening of the relationship between mRNA and lncRNA’ in the supplementary material.

By integrating the very comprehensive MTIs and LTIs mentioned above, we defined the relationship between gene and noncoding RNAs as ‘strong’ and ‘weak’ according to their experimentally validated means. Specifically, we defined the relationship as *strong* after verification by multiple powerful verification methods, such as Luciferase reporter assay or Clip-seq. The relationship verified by other experimental methods, such as qRT-PCR/Western blot, were deemed as insufficient evidence and the predicted relationship called *weak*.

### Pathway databases and noncoding RNAs set enrichment analysis

In this update, a total of 222 human pathways were manually redrawn by integrating information from non-coding RNAs and original KEGG pathways, which cover five main categories in humans, specifically: genetic information processing, environmental information processing, cellular processes, organismal systems, and human diseases. To determine whether specific noncoding RNAs are associated with specific signaling pathways, we used enrichment analysis functions. The enrichment significance P-value for that reference set is calculated as:

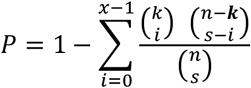

For each pair of noncoding RNA and signaling pathway, we applied a hypergeometric test (64) to evaluate whether the pathway contains significantly more target genes than expected by chance. The p-values were BH-adjusted and a significance level of 0.05 was selected.

### Web server implementation

The current version of NcPath was organized using MySQL 5.7.17 (http://www.mysql.com) and operates on a Linux-based Aliyun Web server. Adopting the idea of separating front and back ends, the web server uses Golang Gin v1.7.7 and Python V3.6 mixed development. NcPath was built using microservice architecture, and gRPC was used for communication between microservices. The frontend is built using common HTML, CSS and Javascript libraries, including the Vuetify framework for styling, Apache Echarts for the visualization library, and the Vue.js framework for the side. Vue.js is a popular front-end technology and the mainstream progressive framework for building user interfaces. We chose this software due to better performance advantages compared to other front-end mainstream frameworks.

## RESULTS

### Overview of the NcPath database

The main elements of NcPath, including the redrawn KEGG pathways, the collection of MTIs/LTIs/ceRNA networks, and the user interface are shown in Figure 1. NcPath provides a user-friendly interface to browse, search and download detailed information on all reference noncoding RNAs sets in pathways. In particular, NcPath provides enrichment analysis of noncoding RNA sets.

### Browsing interface for conveniently retrieving noncoding RNA sets in pathways

To provide a user-friendly interface, we redesigned the displayed pathway map based on KEGG datasets. NcPath enables users to intuitively determine the non-coding RNAs that target a gene in a pathway, the genes that are regulated by non-coding RNAs in a pathway, and the non-coding RNAs that participate in a pathway (Figure 2A). The user can efficiently browse the interactions of interest with the pathway-centric, miRNA-centric and the lncRNA-centric, in one-to-many or many-to-many relationships. In the choose box in the upper left corner of the ‘Browse’ interface, users can select or search for a specific pathway by inputting the pathway’s name and click on the ‘GO’ button on the right to enter the pathway screen. When users browse a pathway map, the red number shown in the upper right corner of the gene represents the sum of the number of noncoding RNAs (miRNAs and lncRNAs) known to regulate this gene (Figure 2A). The number of relationships between the gene and the associated noncoding RNAs will be displayed in the floating window when the cursor of the mouse touches the region where the gene is located in the map (Figure 2A). The right column shows the list of noncoding RNAs related to target gene of choice. There are two main categories, including miRNAs and lncRNAs, each with a list of subcategories with strong and weak relationships. If users wish to obtain the gene set related to the noncoding RNA of interest in reverse, they can place the mouse cursor on the RNA entry in the list on the right column, and all genes related to the target noncoding RNA will be displayed with an orange border. In addition, there is a search box in the middle of the screen that allows users to directly locate the input target gene or the related genes of input non-coding RNA. The specific details on the gene and related noncoding RNA in the new pathway map can be viewed upon clicking. For example, the user can click on the RNA located in the right column to enter its basic annotation information and obtain the list of all pathways associated with this RNA (Figure 2E). If the user clicks the gene icon of the pathway map, the page will redirect to the interactive network and annotation interface (Figure 2B,2C). The interactive network diagram (Figure 2B) allows users to visualize the interaction relationships between noncoding RNAs and protein-coding genes. NcPath provides five types of interactions based on experimental validations and predictions of datasets, including miRNA-strong, lncRNA-strong, miRNA-weak, lncRNA-weak and ceRNA, which are represented by five different colors in the network (Figure 2B). The information being displayed can be controlled by the user by clicking on the relationship icons. In addition, the MTIs/ LTIs references related to the target gene will be displayed in the middle of the page, including the PMID hyperlink, the support type and the method used for experimental verification of the interactions (Figure 2C). To facilitate further studies of the function and mechanism of noncoding RNAs, ceRNA networks are also displayed in the list at the bottom of the Results page.

**Figure 2.**
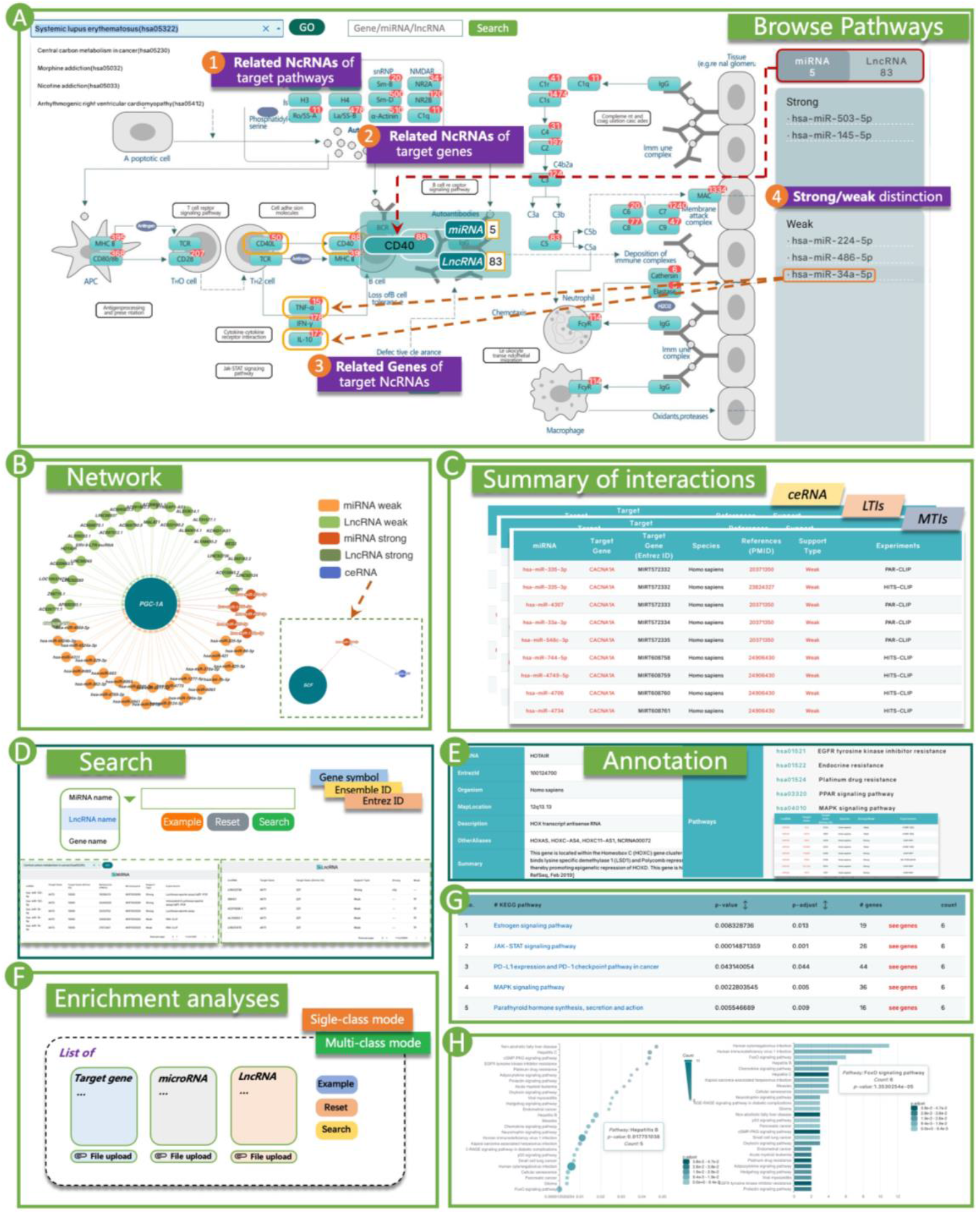
Introduction and usage of NcPath. (A) By browsing the pathway ‘Systemic lupus erythematosus’, users can intuitively obtain (i) Related ncRNAs of target pathways; (ii) Related ncRNAs of each target gene; (iii) Related Genes of target ncRNAs; (iv) Strong/weak distinction. (B) The noncoding-related interaction network of the gene PGC-1A and the ceRNA network for the gene SCF. (C) The table summarizing the interactions of the gene PGC-1A in the pathway ‘Systemic lupus erythematosus’, including MTIs, LTIs and ceRNA information. (D) Users can search by gene and noncoding RNA to obtain comprehensive information, and intuitively browse MTLs, LTIs and ceRNA networks related to the target pathway. (E) The display containing annotation information, associated pathways and noncoding RNAs-target interaction tables from the search function. (F) NcPath provides two input modes of enrichment analysis, including the single-class and multi-class modes. (G) Table with the results of the enrichment analysis. (H) Bubble results and column charts of enrichment analysis.

### Effective online tool for noncoding RNA set enrichment analysis

After generating the noncoding RNA and target gene network with NcPath, users can proceed to the interpretation of the results, an essential part of the analysis process. NcPath provides noncoding RNA (lncRNA and miRNA) set enrichment analysis for users that includes a total of two input modes (Figure 2F). In the single-class mode, the user can decide from five upload options, i.e., whether to select a single lncRNA, a single miRNA, a list of either lncRNAs, miRNAs or genes. In addition, a multi-class mode query (lncRNAs, miRNAs and mRNA are included) can be initiated to find function pathways that have a significant impact on these merged inputs. The details on the enrichment results are shown on the return page; users can click on the title bar of the table to sort, or on the ‘p-adjust’ to use the adjusted P-value and preferentially display the pathway information that is significantly enriched for the target noncoding RNAs set (Figure 2G). The bottom part of the page shows the visualization results of the enrichment analysis, including bubble charts and histograms (Figure 2H). Users can click the pathway name in the enrichment result list to enter the pathway for browsing, or click the number in the ‘genes’ column to obtain the gene set enriched in the pathway. All significant reference pathways and visualization results for the enrichment analysis are provided for review and download. NcPath thus provides rapid access to commonly used gene/miRNA/ lncRNA set enrichment tools that help researchers focusing on the relevant hits.

### Newly designed and more user-friendly interface

The ‘Search’ page is organized as an interactive table that allows users to quickly search for noncoding RNA sets and pathway-related information. Users can search for noncoding RNAs of interest using multiple features, including gene symbol, Entrez ID or Accession (Figure 2D). The results page of the query returns the basic information and related pathways of noncoding RNAs or genes (Figure 2E). For example, if users enter a lncRNA search, the program displays a reference table for each lncRNA that includes gene symbol, Entrez ID, organism, map location, description, other aliases, and summary. This information enables a rapid understanding of the related functions of the specific lncRNA. Users can browse more information from the drop-down menu on the right. In particular, the results page returns all pathways related to the query noncoding RNAs, which are available to users upon clicking on each category. It is worth mentioning that, in the process of browsing information, users can enter the specific annotation webpage when encountering nouns of interest in the pathway map or annotation information through the hyperlinks. This will allow users to obtain more relevant details, e.g., by selecting the hyperlink in the references of the noncoding RNA to browse in more detail. In the lower part of the ‘Search’ interface, users can browse all noncoding RNA-related interaction tables related to the target pathway by searching the pathway name, which makes it easier to directly obtain the desired information (Figure 2D).

### Data download

The ‘Download’ page was organized as an interactive table. All reference sets of the pathways and noncoding RNAs were arranged and sorted into separate files and are available for download in our database (.txt format). Users can download the reference collection as valuable supplementary data to perform in-depth experimental studies.

### Use case 1––noncoding RNAs related to critical illness in COVID-19

As the first use case, we studied the target genes and pathway networks of CDKN2B-AS1, a gene that has been previously described in humans and associated with various functions [87,88]. CDKN2B-AS1 is involved in a large number of pathways, as shown in Figure 3A, including ‘Human immunodeficiency virus 1 infection’ and ‘coronavirus disease-COVID-19’. For example, based on the research of coronavirus disease-COVID-19, we were able to search for this gene in the NcPath database to obtain the total interaction set of all noncoding RNAs involved in the target pathway (Figure 3B). We also browsed the gene network in the pathway map and the regulation of each gene by noncoding RNAs (Figure 3C, D). The related-search functionality highlights interactions between CDKN2B-AS1 and 8 target genes (Figure 3E) in this pathway, including IL6 and OAS3. The OAS gene cluster is not only affected by other genes, but also regulated by 35 miRNAs and 50 lncRNAs, and there is an interaction relationship between lncRNA-RMRP and OAS1 (Figure 3C, F). In addition, we found that IFNAR2 is regulated by 163 miRNAs and 16 lncRNAs (Figure 3F). Previous research showed that low expression levels of IFNAR2 are associated with life-threatening COVID-19 disease [89], and that the interferon-inducible oligoadenylate synthetase (OAS) genes are implicated in susceptibility to SARS-CoV, based on candidate gene association studies performed in Vietnam and China [90,91]. The lncRNA CDKN2B-AS1 and other noncoding RNAs can thus be used as potential biomarkers for COVID-19 in future studies. Our results show that NcPath’s interactive regulatory network can aid the rapid discovery of potential regulatory relationships, and help guiding biological experiments.

**Figure 3.**
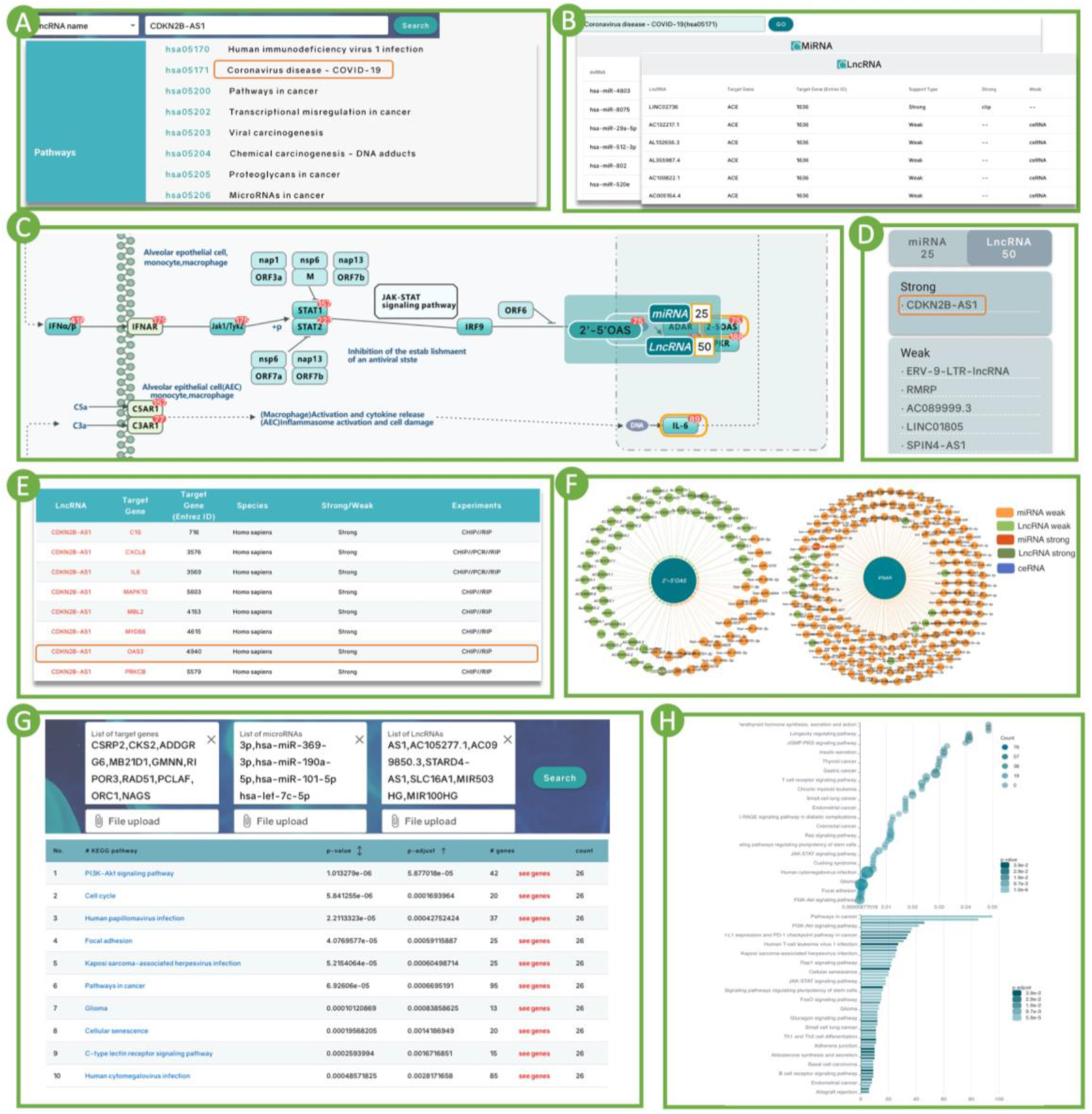
Use cases of NcPath. (A) The related pathways of target noncoding RNA (lncRNA CDKN2B-AS1). (B) The sets of MTIs and LTIs with information about coronavirus disease-COVID-19. (C) Partial display of the interactive diagram of coronavirus disease-COVID-19. (D) The list of lncRNAs and miRNAs that interact with the OSA gene cluster. (E) The summary of lncRNA-target interactions table of CDKN2B-AS1 in coronavirus disease-COVID-19. (F) The interaction networks of the OSA gene cluster and IFNAR2. (G) Results table for enrichment analysis of the squamous cell carcinoma. (H) Bubble and column charts showing enrichment analysis results.

### Use case 2––integrated lncRNA/miRNA/gene analysis in squamous cell carcinoma

The regulation of gene expression in physiological and pathophysiological processes is a central question in biomedical and life science research. To explore the role played by differential miRNAs, lncRNAs and mRNA expression patterns simultaneously, it is necessary to implement enrichment analysis of these genes and noncoding RNAs. We used NcPath to perform functional analyses on the genes differentially expressed in tongue squamous cell carcinoma [92]. The dysregulated mRNA, miRNA, and lncRNA expression profiles of squamous cell carcinoma and normal tissues were extracted from The Cancer Genome Atlas (TCGA). The 10 most significant pathways uncovered are provided as a results table (Figure 3G), with the bubble chart and histogram shown in Figure 3H. The top hit with an adjusted P-value of 5 × 10-5 was the ‘PI3K-Akt-signaling pathway’, and a previous study showed that miR-22 regulates this pathway in tongue squamous cell carcinoma [93]. The second most significant hit was ‘cell cycle’, with an adjusted P-value of 1.6 × 10-4, followed by ‘human papillomavirus infection’, and ‘focal adhesion’. Importantly, numerous experiments demonstrated these pathways affect the occurrence and development of tongue squamous cell carcinoma [94-98]. This use case again demonstrates the practical application of NcPath, which allows for comprehensive analysis using pairings of three RNA classes, and helps researchers focusing on potentially more relevant biological findings.

## CONCLUSIONS AND EXPECTATIONS

We present a significant update of our web server NcPath that allows for the integrative analysis of miRNA, lncRNA and target genes with pathway interaction networks. While the original version was focused on miRNA, we now offer support for lncRNA and a higher number of pathways. NcPath is the first database providing an enrichment analysis on miRNAs, lncRNAs and mRNAs together as input. The development of new technologies and the accumulation of experimental data, resulted in the generation of an increasing number of miRNA-, lncRNA- and ceRNA-realated information. We will also include additional experimental sets to extend our data sources and support more powerful enrichment analysis tools. In the future, Ncpath will continue supplementing more non-coding RNA classes and new features to further enhance functional interpretations, following the developments in biology. In addition, we will focus on expanding the number of species and collections, and providing users with more efficient enrichment analysis methods in the future.

## Supporting information

Screening of the relationship between mRNA and lncRNA

## DATA AVAILABILITY

All the data could be downloaded from http://ncpath.pianlab.cn/.

## SUPPLEMENTARY DATA

Supplementary Data are available at NAR Online.

## FUNDING

The Fundamental Research Funds for the Central Universities (No. JCQY202108) and Startup Foundation for Advanced Talents at Nanjing Agricultural University (No. 050/804009).

## Notes

### Competing Interest Statement

The authors have declared no competing interest.

