## Supplementary material for "NcPath: A novel tool for visualization and enrichment analysis of human non-coding RNA and KEGG signaling pathways": Screening of the relationship between mRNA and lncRNA

### The distribution of lncRNA-mRNA interaction scores:

10,000 scores of lncRNA-mRNA interaction were randomly selected from the five interaction categories of Co-expression, ceRNA (number of shared miRNAs), CO-TF, Co-HM and LncPro score respectively, and their probability density distribution plots were shown as follows:

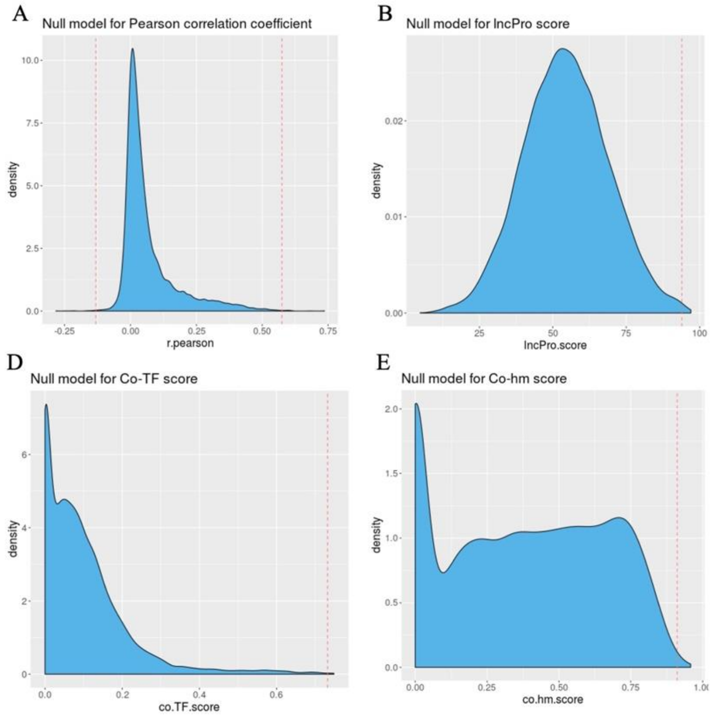


**Figure S1. The probability density distribution plots of five interaction categories.** (A-E) The probability density distribution plots of Co-expression, LncPro score, ceRNA, CO-TF score and Co-HM score.

### Reliable lncRNA-mRNA interaction pairs:

For the above five kinds of interaction relations, Monte Carlo simulation was used to evaluate whether a lncRNA-mRNA pair was significantly reliable. We chose 10, 000 scores randomly each time to the cutoff score of P-value=0.001. This process was repeated 10, 000 times and the average cutoff value is finally determined (see the red line in the figures above, and the table below). Bilateral test was used for co-expression, and the other 4 lncRNA-mRNA interaction relations were tested on the right side.

**Table S1. The average cutoff values of five interaction relations.**

| **Interaction relations** | TF | Hm | Cor | Lncpro | ceRNA |
|---|---|---|---|---|---|
| **Cutoff** | 0.7322795 | 0.9114914 | 0.5748191<br>-0.1311137 | 93.93853 | 50 |

With the above cutoff as the boundary, the lncRNA-mRNA pairs with significant scores in the five interaction relationships were considered to be reliable, that is, the authentic relationship pairs of lncRNA-mRNA interaction in each interaction category. We finally obtained seven types of reliable interactions, including the five relationships mentioned above, cis-regulation interactions and clip-validated interactions. Then the overlap between the seven types of reliable lncRNA-PCG interactions can be seen in the following figure.

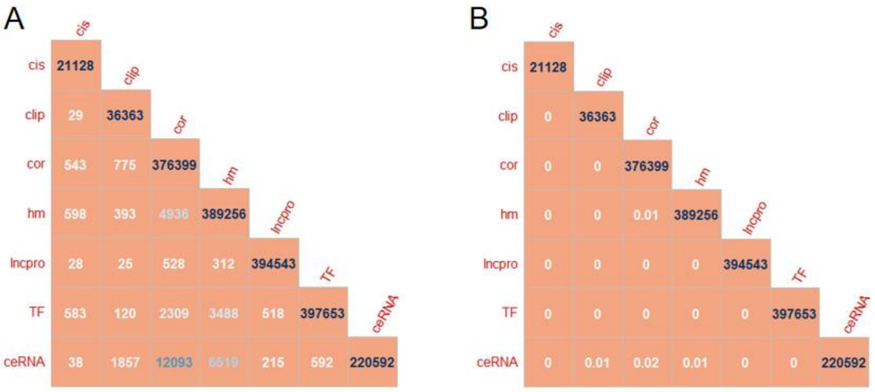


**Figure S2. Then the overlap between the seven types of reliable lncRNA-PCG interactions.**
(A)The overlap number between reliable lncRNA-PCG interactions derived from 7 methods.
(B)The similarity between reliable lncRNA-PCG interactions derived from 7 methods.

The dark blue number on the main diagonal represents the number of reliable lncRNA-PCG interaction pairs obtained by each method. The similarity in Figure B is defined as:

$$\text{similarity}(S,\ T) = \frac{|S \cap T|}{|S \cup T|}$$

S and T represent the lncRNA-PCG interaction pair sets obtained by two different methods respectively.

The above results showed that the reliable lncRNA-PCG interaction pairs obtained by different methods had almost no overlap (the highest similarity between the ceRNA and co-expression was 2%), that is, the reliable lncRNA-PCG interaction obtained from 7 different methods had a relatively large difference.
A total of 1800651 reliable lncRNA-PCG interaction pairs from 7 methods were identified. Then, we observed the number distribution of PCGs interacting with each lncRNA using the reliable interaction pairs (see **Figure S3**). Most lncrnas had reliable interaction with a small number of PCGs, and a few lncRNAs had reliable interaction with a large number of PCGs. There were 2722 lncRNAs that had no reliable interaction with any PCGs (this part of lncRNAs can be considered as having weak regulatory function), accounting for 15% of the total number of lncRNAs. There are 2879 lncRNAs with more than 100 reliable interaction mRNAs, accounting for 16% of the total lncRNAs, which may have strong regulatory function.

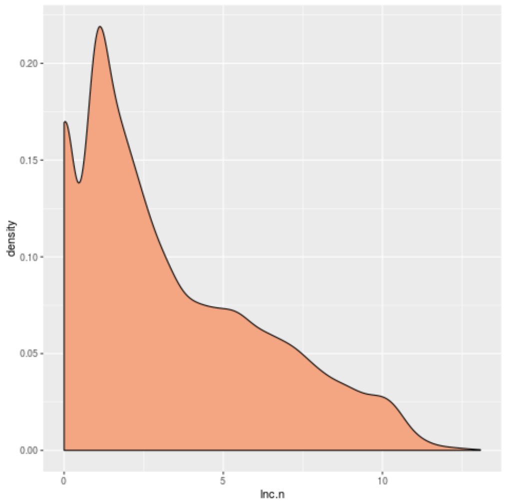


**Figure S3. The number distribution of PCGs interacting with each lncRNA using the reliable interaction pairs.**
